# Rule-dependent transformation of observed actions into self-actions in macaque ventral premotor cortex

**DOI:** 10.64898/2026.07.24.740118

**Authors:** Lilei Peng, Alexander Lappe, Shengjun Wen, Silvia Spadacenta, Martin Giese, Peter Thier, Jörn K. Pomper

## Abstract

The ventral premotor cortex (PMv) has been implicated in both action selection and the perception of observed actions, but it remains unclear whether PMv activity during action observation reflects the observed action itself, or variables related to the observer’s own action when the observed action becomes behaviourally relevant. Here, we recorded neural activity in macaque PMv during a task that dissociated observed action, rule context, and the subsequently selected self-action. Population activity during observation was already biased toward the upcoming self-action and became increasingly aligned with it. At the level of single neurons, subsets showed modulation by rule and required self-action beyond the observed action. Task-related variables coexisted during observation, rather than being organized into distinct sequential stages. These findings indicate that PMv activity during action observation is not solely determined by the observed action, but instead reflects variables related to the selection of the agent’s own action, consistent with a transformation from observed action to self-action.

## Introduction

The premotor cortex has been studied from two dominant perspectives. The first perspective considers it a key site for the selection and implementation of goal-directed actions^1,2^. Within the frameworks of action selection, including the affordance competition framework, multiple potential actions can be represented in parallel and compete for execution, with environmental information incorporated into these representations insofar as it is relevant for the agent’s behaviour^3–5^. The second perspective posits a role for ventral premotor cortex (PMv) in the perception of others’ actions, following the discovery of mirror neurons whose activity is modulated during both action execution and observation^6–8^.

Extensive work has been carried out guided by these two concepts. Yet, it remains unclear whether and to what extent these perspectives can be integrated, rather than being viewed as largely incompatible. More specifically, it remains unclear how the two perspectives relate when observed actions become directly relevant for the observer’s own behaviour^4,8,9^. In particular, it is not well understood whether representations of observed actions are maintained as such, rather than reflecting variables relevant for the observer’s own action decisions. Resolving this question is critical for understanding how action observation contributes to behaviour.

In an attempt to find an answer, we investigated PMv activity in rhesus monkeys in a task that explicitly dissociated an observed action, a rule defining a particular context, and the subsequent action selection. Monkeys first observed an object manipulation (twist or lift) and then, depending on the previously presented rule (same or other), selected which action to execute themselves. This design allowed us to determine whether PMv activity reflects the observed action, the derived self-action, or the context provided by the rule, and how these components evolve over time.

From the perspective of action selection frameworks, including the affordance competition framework^5^, an observed action, if treated as an environmental variable, may be rapidly incorporated into variables related to the observer’s own action. Under this view, PMv activity is expected to increasingly reflect the selected action as it becomes determined. In contrast, an action-perceptual account predicts that PMv initially maintains a representation of the observed action during viewing that remains dissociated from subsequent action selection^8^.

Our results show that PMv activity during action observation reflects not only the observed action, but also task-related information relevant for the selection of the upcoming self-action, suggesting that PMv activity during action observation may already be related to an ongoing action selection process.

## Results

We recorded neural activity in ventral premotor cortex (PMv) as monkeys performed a task in which they observed one of two object-directed actions – lifting (‘Lift’) or twisting (‘Twist’) an object – and subsequently executed one of the two actions depending on the prevailing rule (Fig. 1). Each trial consisted of four phases. During the cue phase, a visual rule cue indicated whether the monkey should perform the same or the alternative action relative to a subsequently observed action. Rule identity was conveyed by cue color and reinforced by a brief vertical displacement of the cue (10° upward or downward). The monkey was expected to follow the displaced cue with its gaze, but the fixation window (±9° visual angle) allowed some flexibility in gaze position. During the observation phase, the monkey viewed a video of another monkey performing one of the two possible actions. After a delay period (preparation phase), a go signal instructed the monkey to execute the required action (execution phase).

**Figure 1.**
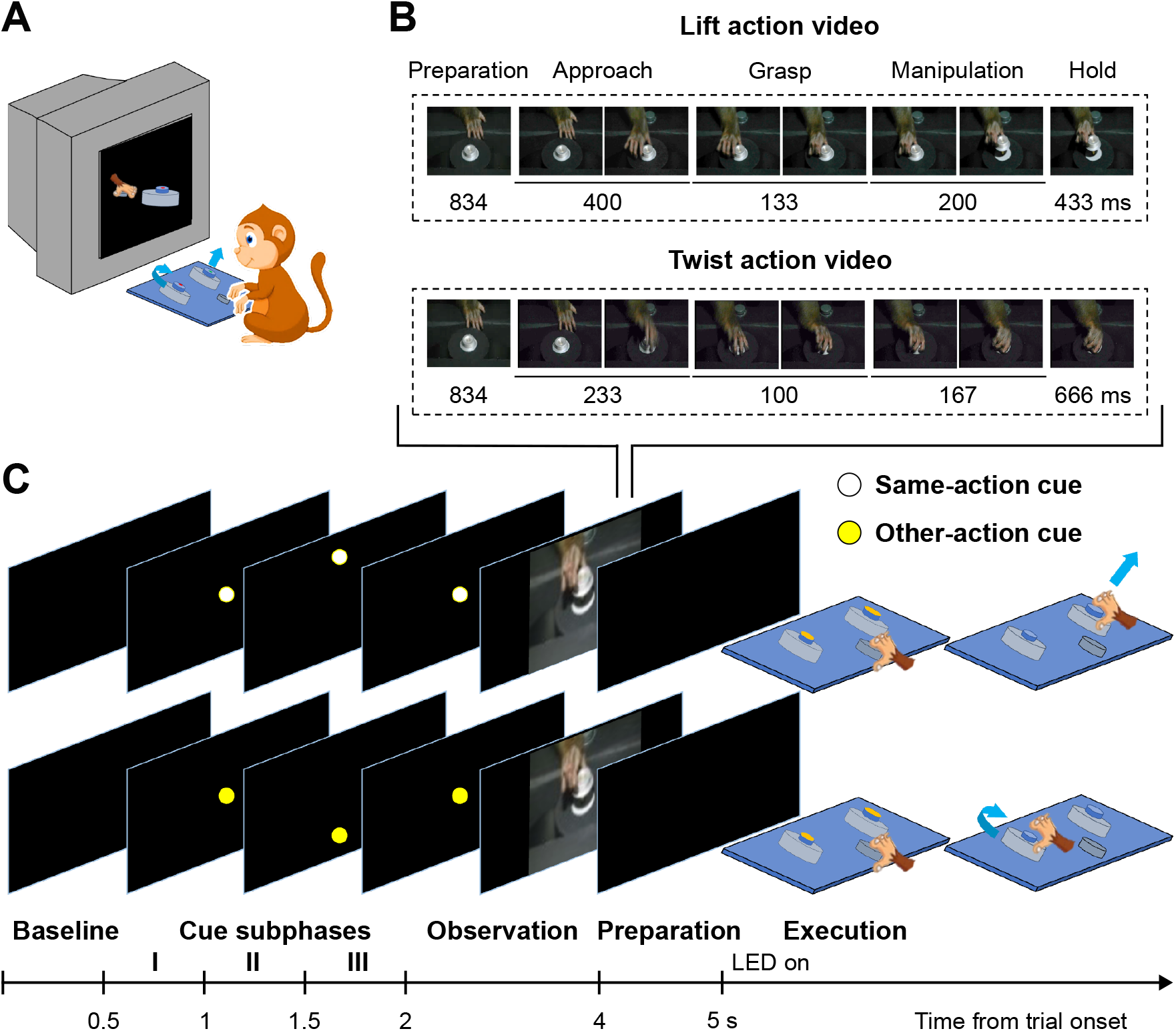
Experimental paradigm. (A) Schematic of the task. (B) Two types of video clips were presented (lift and twist actions; 2 s duration), subdivided into five epochs: preparation, approach, grasp, manipulation, and hold. (C) Example trial (lift video shown). Trials began with a baseline phase (0–0.5 s), followed by a three-part cue phase (0.5–2 s) in which a coloured dot (white or yellow) indicated the rule condition. The dot remained at the centre (Cue I), shifted vertically (Cue II; ±10° visual angle), and returned to the centre (Cue III). The colour of the dot specified the rule: a same-action cue (white) instructed the monkey to perform the observed action, whereas an other-action cue (yellow) instructed the monkey to perform the alternative action. During the observation phase (2–4 s), an action video was presented, followed by a preparation phase (4–5 s). The simultaneous illumination of LEDs on top of both objects served as the go signal, after which the monkey released the home button and executed the instructed action. Correct performance was rewarded with a drop of water. Intervals are not drawn to scale.

The task was performed in two paradigms that differed in how rule and observed action information were distributed across trials (Fig. 2). In the video-blocked paradigm, the observed action was fixed within a block, whereas the rule cue (same vs. other) varied randomly across trials. In the cue-blocked paradigm, the rule cue was fixed within a block, whereas the observed action varied randomly across trials.

**Figure 2.**
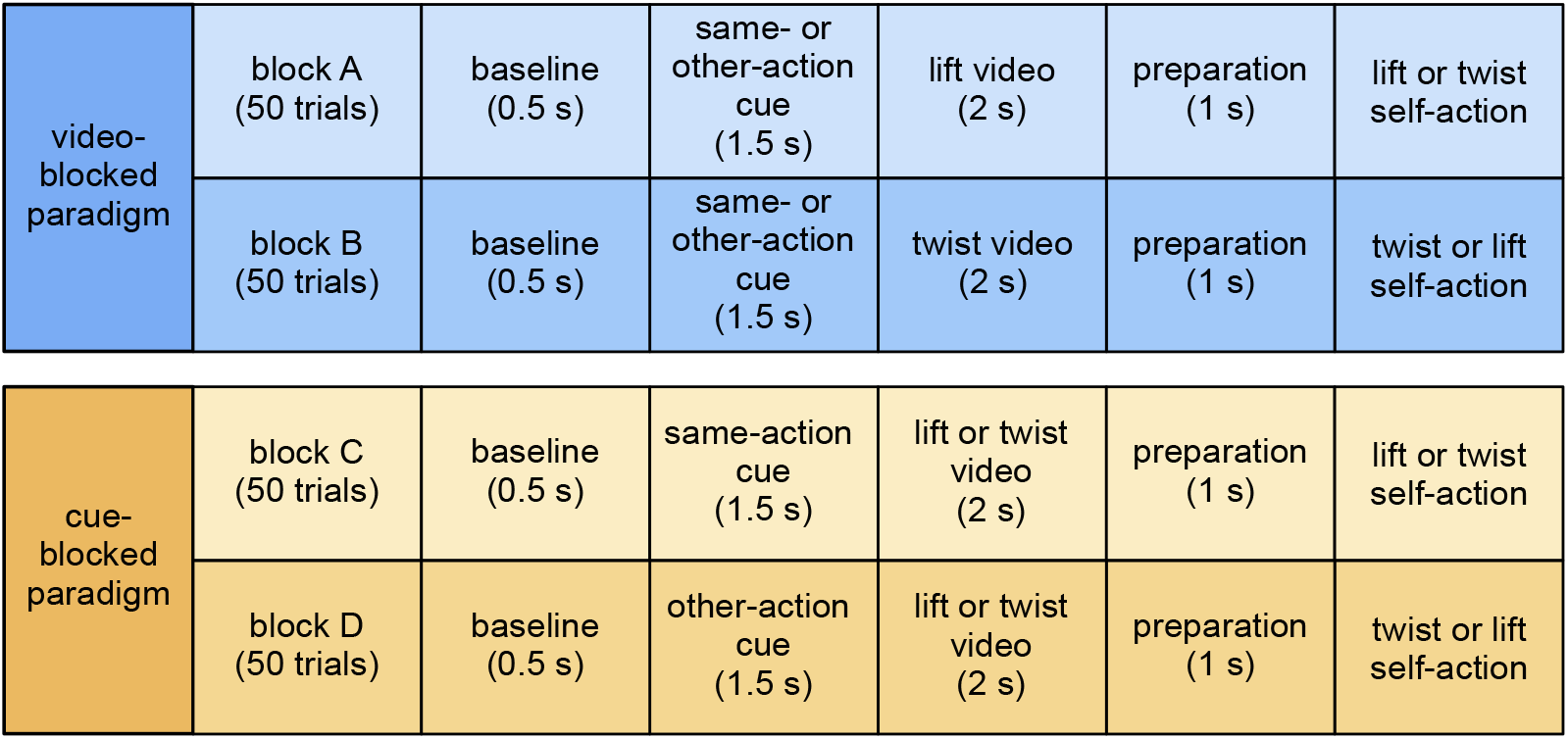
Block structure of the two task paradigms. In the video-blocked paradigm (top), the observed action (lift or twist) remained constant within a block, whereas the rule cue varied across trials. In the cue-blocked paradigm (bottom), the rule cue remained constant within a block, whereas the observed action varied across trials. Each trial followed the same sequence of phases (see Fig. 1C).

We recorded 859 neurons in the video-blocked paradigm and 288 neurons in the cue-blocked paradigm (from two monkeys). Of these, 500 (58.2%) and 228 (79.2%), respectively, were classified as task-related based on their activity during observation and execution (see Methods) and included in subsequent analyses. This selection focused the analysis on neurons relevant for linking action observation to self-action.

To determine whether PMv population activity reflected the rule context, the observed action, or the subsequently selected self-action, and how these representations evolved over time, we developed a joint PCA tailored to the structure of our task. Standard PCA captures dominant patterns of variance in population activity but does not distinguish, how this variance relates to specific task variables. Our approach extends PCA to account for the block structure of the task, which does not permit a conventional demixed decomposition (e.g., demixed PCA^10^).

To avoid confounds from non-specific differences between blocks, we focused on differences in activity between conditions within each block. This allowed us to compare the representation of the selected self-action against the task variable manipulated within each block (rule in the video-blocked paradigm and observed action in the cue-blocked paradigm). The joint PCA identifies components in this differential activity between conditions that capture variance shared across both blocks and classifies them according to how this shared structure aligns across blocks (see Methods). Specifically, components were defined such that their projections aligned across blocks either for conditions requiring the same self-action or according to the task variable (i.e., rule or observed action). This classification defines how shared population structure is interpreted in the component projections shown in Figs. 3 and 4.

**Figure 3.**
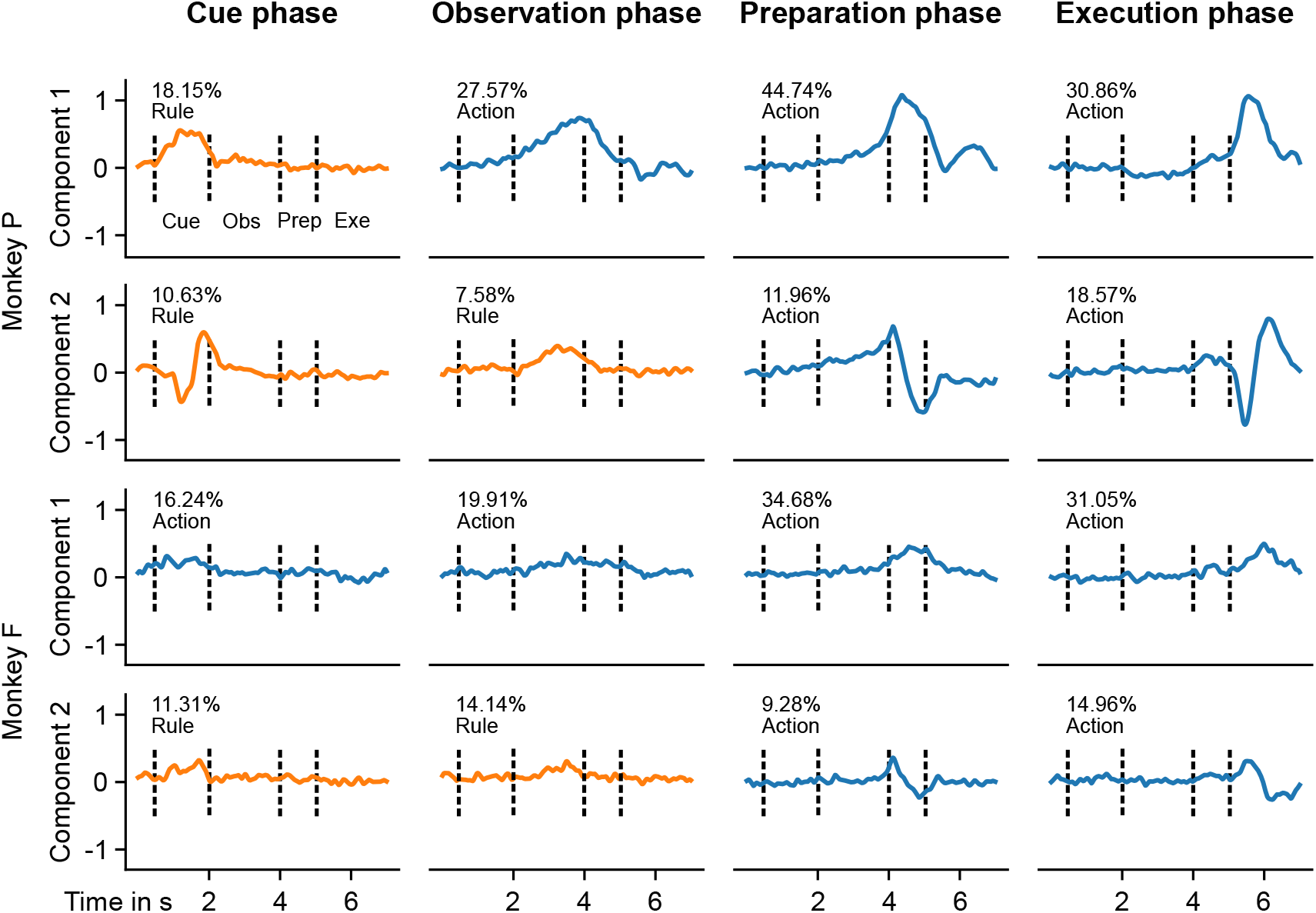
Joint principal components for the video-blocked paradigm. Each panel shows the time course of population activity projected onto a single component. Components were computed separately for each trial phase (cue, observation, preparation, execution; columns), but projections are shown across the full trial. Components define directions in population activity that capture dominant patterns of variance across neurons. Each component is classified according to how its projections align across blocks, resulting in projections that are organized either by self-action (blue) or by rule cue (orange), as indicated in each panel. The vertical axis reflects the projection of population activity onto the component over time. Because the sign of the projection is arbitrary, its absolute direction (positive or negative) is not meaningful. Instead, interpretation focuses on the magnitude and temporal profile of the signal, indicating when the component is active during the trial. Dashed lines indicate the onset of the cue, observation, preparation and execution phases. Text indicates the proportion of variance explained by each component. Data from two monkeys are shown (P, top; F, bottom).

**Figure 4.**
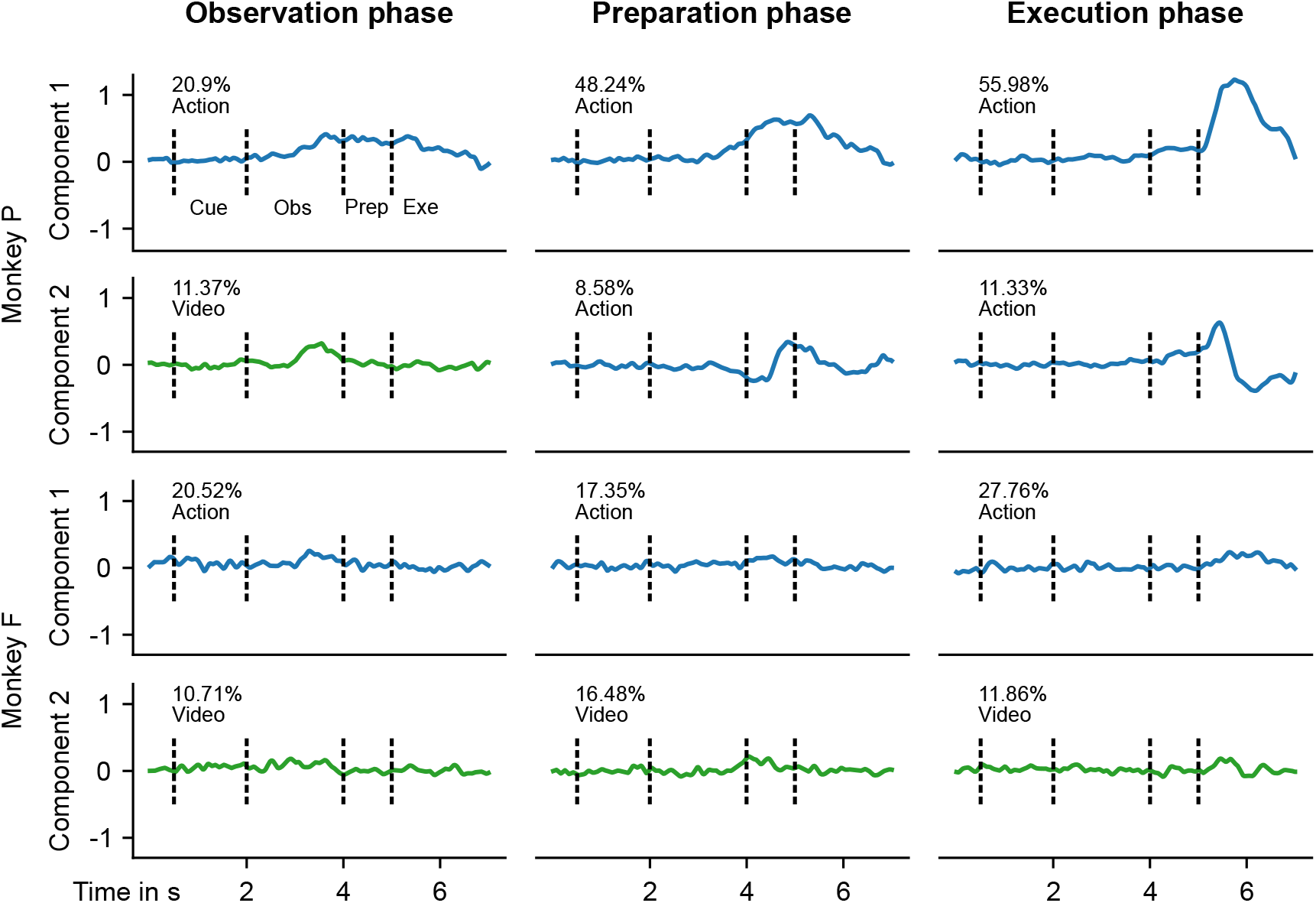
Joint principal components for the cue-blocked paradigm. Same format as in Fig. 3. Components are aligned either with executed action (blue) or observed action (green), as indicated in each panel.

We first examined how population activity evolves over time with respect to the task variable and the subsequently selected self-action. In the video-blocked paradigm, projections onto rule-aligned components showed strong modulation during the cue phase, consistent with the availability of rule information prior to action observation (Fig. 3). During the observation phase, however, projections onto self-action-aligned components emerged and increased in magnitude, exceeding those of rule-aligned components. A similar pattern was observed in the cue-blocked paradigm, where projections onto both observed-action-and self-action-aligned components were present during observation, with self-action-related signals already clearly visible at this stage (Fig. 4). Across both paradigms, this coexistence was followed by a pronounced shift during the preparation and execution phases, in which projections onto self-action-aligned components became dominant, while projections onto task-variable-aligned components diminished. This transition was observed in both monkeys, although the strength and temporal sharpness of the effect varied across individuals.

This shift in population structure was further quantified by the proportion of shared variance explained by the first four components aligned with each variable. In the video-blocked paradigm, self-action-aligned components accounted for approximately 50–60% of the shared variance during preparation and execution in both monkeys, whereas rule-aligned components contributed substantially less (Table 1). A similar pattern was observed in the cue-blocked paradigm, with self-action-aligned components dominating in later phases and reaching up to ∼75% of the shared variance in Monkey P, while remaining lower but still predominant in Monkey F (Table 2).

**Table 1.** Proportion of variance explained by the first four components in the video-blocked paradigm.

| Phase | Monkey P |  | Monkey F |  |
| --- | --- | --- | --- | --- |
|  | Self-action | Rule | Self-action | Rule |
| Cue | 8.3% | 35.9% | 22.7% | 17.4% |
| Observation | 27.6% | 16.9% | 28.6% | 23.0% |
| Preparation | 56.7% | 6.7% | 49.8% | 8.5% |
| Execution | 59.5% | 1.6% | 52.7% | 7.5% |

**Table 2.** Proportion of variance explained by the first four components in the cue-blocked paradigm.

| Phase | Monkey P |  | Monkey F |  |
| --- | --- | --- | --- | --- |
|  | Self-action | Video | Self-action | Video |
| Observation | 28.5% | 18.5% | 28.5% | 20.5% |
| Preparation | 60.5% | 5.8% | 26.2% | 29.8% |
| Execution | 75.7% | 0.0% | 36.4% | 20.3% |

Together, these results indicate that PMv population activity does not maintain a stable representation of the observed action or rule context throughout the trial. Instead, task-related information present during cueing and action observation is progressively reorganized, resulting in a population structure aligned with the selected self-action, which dominates during movement preparation and execution.

To assess how population-level effects relate to single-neuron responses, we fitted generalized linear models to spike counts in the video-blocked paradigm, using the observed action as a baseline model and testing for additional modulation by rule and required self-action.

During the observation phase, a subset of neurons exhibited responses that could not be explained by the observed action alone. Specifically, the model based on rule and/or self-action provided a significantly better fit for 21.4% of neurons in monkey P (81/379) and 10.7% in monkey F (13/121), indicating modulation beyond the observed action.

We next examined the structure of this modulation by analyzing the corresponding regression coefficients. Neurons were classified according to whether their responses were best described by self-action, rule, or an interaction between the two (Table 3). This analysis revealed a heterogeneous distribution of tuning profiles across neurons. In both monkeys, subsets of neurons were selective for the required self-action, independent of the rule, while others exhibited sensitivity to the rule or to specific combinations of rule and required self-action.

**Table 3.** Proportion of neurons classified as tuned to self-action, rule, or their interaction, among neurons whose responses were significantly better explained by a model including rule and/or required self-action than by a model based on the observed action alone. Percentages are relative to this subset, separately for each phase and monkey.

|  | Self-action |  | Rule |  | Interaction |  |
| --- | --- | --- | --- | --- | --- | --- |
|  | Monkey P | Monkey F | Monkey P | Monkey F | Monkey P | Monkey F |
| Cue phase | 25.9% | 44.4% | 57.4% | 0 | 16.7% | 55.6% |
| Observation phase | 70.4% | 30.8% | 17.3% | 46.2% | 12.3% | 23.1% |

A similar heterogeneity of tuning profiles was already present during the cue phase (Table 3), indicating that task-related information is reflected at the level of single neurons prior to action observation. Across both phases, these results show that modulation beyond the observed action is not uniformly expressed across neurons, but instead distributed across subpopulations with distinct response profiles. This pattern at the level of single neurons is consistent with the population-level structure revealed by the joint PCA, in which activity reflects task-related variables during early phases and tends to become aligned with the selected self-action over time, although the strength of this effect varied across monkeys. These findings indicate that neural responses in PMv during action observation are not solely determined by the observed action, but already reflect task-related information relevant for the upcoming action.

## Discussion

We asked whether PMv activity during action observation reflects the observed action itself or instead relates to the selection of the observer’s own action when the observed action is informative for the action to be executed. Our results support the latter, showing that PMv activity during observation already reflects the emerging selection of the upcoming self-action. This is evident at the population level, whose activity progressively aligns with the selected self-action, and at the level of single neurons, with a subset of neurons exhibiting modulation by rule and required self-action beyond the observed action.

Our findings place PMv activity during action observation within a framework of action selection and sensorimotor transformation. Previous work has demonstrated population-level transformations from visual inputs to representations of the agent’s own upcoming actions^11^. Here, we extend this work by showing that similar transformations occur when the relevant visual input consists of observed actions that inform the observer’s own behaviour. Consistent with this view, PMv population activity during observation was already biased toward the subsequently executed action and modulated by rule context, indicating that observed actions are rapidly integrated into the evolving action selection process. This interpretation is in line with studies showing that premotor and parietal populations represent potential actions in a distributed manner and that activity progressively converges onto the selected action over time^12,4,13^.

This interpretation does not preclude that PMv activity during action observation can, in other contexts, reflect a representation of the observed action itself^8^. Previous studies in premotor cortex, including PMv and PMd, have reported mixed evidence for shared representations during action execution and observation, with little overlap at the level of single neurons but partial similarity at the population level^14–19^. Our results are compatible with these findings, but suggest that such activity should not be interpreted as necessarily reflecting the observed action per se. Instead, the same population responses may also reflect variables relevant for the selection of the observer’s own action, depending on context^20–23^. In this view, activity during observation does not uniquely specify a representation of the observed action, but can instead participate in a process that links observed information to action selection.

Task-related information was not organized into distinct sequential phases, but instead coexisted during observation, with representations related to rule, observed action, and self-action present in parallel. Over time, population activity showed a gradual shift toward the selected self-action, rather than a discrete transition between perceptual and motor representations, consistent with dynamical systems accounts of population activity. This temporal structure further supports the view that action observation is closely linked to action selection.

From a functional perspective, this organization suggests that observed actions are not maintained as independent representations, but are rapidly transformed into variables relevant for the observer’s own action. Such an organization may facilitate flexible mapping between observed actions and appropriate responses, as required by the task.

The strength of these effects varied across monkeys, particularly in the extent to which activity became dominated by self-action during later phases. This variability may reflect differences in task strategy or in the reliance on specific task variables, but does not alter the overall pattern of results.

A further aspect concerns the relationship between single-neuron tuning and population-level structure. At the level of individual neurons, responses were heterogeneous with different subsets of neurons selective for self-action, rule, or their interaction. At the same time, population activity showed a coherent structure and became increasingly aligned with the selected self-action over time. This suggests that population-level representations do not arise from a homogeneous code across neurons, but instead reflect the contributions of subsets of neurons with different functional roles.

Finally, our results are based on a task in which observed actions are directly linked to subsequent action selection by an explicit rule. Whether similar patterns extend to more naturalistic settings, where such mappings are not constrained by a narrow set of possibilities, remains an open question, particularly in light of recent work showing that premotor representations differ between constrained and freely moving conditions^24^. More generally, our findings suggest that activity in PMv during action observation does not have a fixed functional role, but can reflect how observed information contributes to the selection of the observer’s own action, depending on internal state and environmental context.

## Acknowledgments

We thank Dr. Peter Dicke for the presurgical MRI scans and implant design, assistance with surgeries and postsurgical animal care, and for his major contributions to the development of the experimental setup. We further thank Dr. Friedemann Bunjes for his contributions to the development of the experimental setup, in particular the software for experimental control and data acquisition.

## Author Contributions

L.P., S.W., S.S., P.T., and J.K.P. conceived the study. S.W. trained both monkeys and performed some of the experiments. L.P. performed most of the experiments. A.L. developed the analytical approaches, including the PCA and GLM analyses, and, together with L.P., analyzed the data. M.G. supervised the development of the analytical approaches. L.P. wrote the initial draft. J.K.P. substantially revised and wrote the manuscript. All authors discussed the results and revised the manuscript. P.T. and J.K.P. supervised the project. L.P., A.L., and S.W. contributed equally to this work.

## Methods

### Subjects and ethical approval

Experiments were conducted in two adult male rhesus monkeys (*Macaca mulatta*; Monkey P and Monkey F; 10 and 11 kg). All procedures were performed in accordance with German and European regulations and the National Institutes of Health Guide for the Care and Use of Laboratory Animals. All experimental protocols were approved by the Animal Welfare Commission (§15) of the Regierungspräsidium Tübingen and were supervised by the institutional Animal Welfare Officers of the University of Tübingen and the local veterinary authorities (Landratsamt Tübingen). Veterinary care was provided by the University of Tübingen.

### Surgical procedures

Each animal was implanted with a titanium head post for head fixation and a titanium recording chamber positioned over area F5 in the left hemisphere. Chamber placement was guided by a preoperative anatomical MRI scan. All surgical procedures were performed under aseptic conditions with general anesthesia induced and maintained using isoflurane (0.8–1%) in combination with remifentanil (1–2 µg kg^− 1^ min^− 1^). Vital parameters, including heart rate, core body temperature, blood pressure, and blood gases (PO_2_, PCO_2_), were continuously monitored throughout the procedure. Postoperative analgesia included carprofen (4 mg kg^− 1^, once daily) and buprenorphine (0.01 mg kg^− 1^, 2–3 times daily) and was maintained as required. Animals were allowed to fully recover before resuming experimental procedures.

### Experimental setup

Monkeys were trained to perform an object-directed grasping task based on visual instructions presented on a monitor (Fig. 1A). The right hand was free to move, whereas the left hand was restrained by a baffle.

A custom-built grasping apparatus (“grasping table”) was positioned within reach of the right hand. The apparatus consisted of a plastic plate tilted ∼55° relative to the horizontal and contained three manipulanda: a centrally located home button and two identical cylindrical objects positioned to the left and right at a greater distance from the animal. All manipulanda were equipped with mechanoelectrical sensors to detect initial contact with the object as well as the onset and offset of object manipulation. The two cylindrical objects were visually and haptically identical but required distinct actions. The left object could be lifted vertically (up to 2.6 cm), whereas the right object could be rotated clockwise (∼70°) against a spring force. Both objects were equipped with LEDs at their tops.

Visual stimuli were presented on a 15-inch monitor positioned 44 cm in front of the animal (measured from the eyes). The midline of the head, the home button, and the midpoint between the two objects were aligned and perpendicular to the monitor.

Behaviour was monitored using two infrared cameras, one tracking body movements and one tracking the right eye. Eye position was recorded using an in-house eye-tracking system (50 Hz sampling rate; ∼150 ms latency for online control).

Experimental control and behavioural data acquisition (including eye position) were performed using the open-source software Nrec (developed by F. Bunjes, J. Gukelberger et al.; https://nrec.neurologie.uni-tuebingen.de) running on a Debian Linux system. Electrophysiological signals were acquired via the AlphaLab system (Alpha Omega Engineering).

### Action video stimuli

Visual stimuli consisted of two video clips depicting goal-directed actions (lift and twist) performed by a monkey’s right hand acting on the cylindrical objects used in the task. The actions shown in the videos were performed by one of the two monkeys participating in the experiment. Videos were recorded from a frontal perspective, as if facing the performer.

Original recordings were edited using MATLAB 2018 (MathWorks) to generate the final stimuli. Video clips had a resolution of 700 × 400 pixels and a duration of 2 s (60 frames). When presented on the monitor, they subtended 32° × 18° of visual angle (width × height).

Each video began with the demonstrator’s right hand positioned on the home button, with the target object within reach and the object LED turned off. Both action types (lift and twist) could be subdivided into five epochs (Fig. 1B): preparation, approach, grasp, manipulation, and hold. During the preparation epoch, the hand remained on the home button and pressed it against a spring; this phase was identical across action types. The approach epoch spanned the interval from hand release to object contact, followed by the grasp epoch, defined from initial contact to movement onset. The manipulation epoch extended from movement onset to completion of the action (object reaching its final position), and the hold epoch corresponded to maintaining the object in the final position. Epoch durations differed slightly between lift and twist actions but together spanned the full video duration. Exact epoch durations are provided in Fig. 1B.

### Behavioural task

At the beginning of each experimental session, eye position was calibrated. As the head was fixed via the implanted headpost, eye position corresponded to gaze direction.

Monkeys were required to perform one of two possible actions based on the observation of a video and a task rule. Depending on the rule condition, animals either reproduced the observed action (“same” condition) or executed the alternative action (“other” condition). Thus, each trial was defined by the combination of observed and executed actions, resulting in four conditions: lift–lift (LL), twist–twist (TT), lift–twist (LT), and twist–lift (TL).

A fully randomized design across all conditions was not feasible, as animals were unable to reliably perform the task without stabilizing one task variable over multiple trials. Therefore, two complementary task paradigms were used to dissociate the contributions of rule and observed action. In both paradigms, a rule cue was presented in each trial. In the video-blocked paradigm, the observed action (lift or twist) was fixed within blocks of 50 trials, whereas the rule cue (same or other) varied across trials. In the cue-blocked paradigm, the rule cue was fixed within blocks, whereas the observed action varied across trials (Fig. 2).

Each trial consisted of five phases (Fig. 1C): baseline, cue, observation, preparation, and execution. During the baseline phase (0–0.5 s), the monkey held the home button while the screen remained blank. In the cue phase (0.5–2 s), a coloured dot (white or yellow; radius 1° visual angle) indicated the rule condition. The cue was presented in three successive subphases (each 0.5 s), including a transient vertical displacement (±10°, with direction contingent on colour).

In the observation phase (2–4 s), a lift or twist action video was presented. During the preparation phase (4–5 s), the screen returned to black while the monkey maintained contact with the home button.

The execution phase was initiated by simultaneous illumination of LEDs on both objects (0.5 s), serving as a go signal. The monkey released the home button and performed the instructed action (lift or twist). The action had to be completed within 3.5 s and included maintaining the object in its final position for 1–1.5 s before returning the hand to the home button.

Monkeys were required to maintain fixation within phase-specific fixation windows during the rule-cue and observation phases. During the three rule-cue subphases, fixation windows of ±5°, ±9°, and ±7.5° visual angle were used, with grace periods of 350 ms, 300 ms, and 300 ms, respectively. During the observation phase, fixation windows of ±10° visual angle (±15° in some sessions) with a 100 ms grace period were applied. Trials were aborted if behavioral or fixation requirements were not met. Correct trials were rewarded with a drop of water (0.3 ml). Behavioral and electrophysiological data were recorded simultaneously for offline analysis.

### Electrophysiological recordings

Extracellular action potentials were recorded using commercially available glass-coated tungsten microelectrodes (0.5 or 1 MΩ impedance at 1 kHz; Alpha Omega Engineering). Electrodes were advanced through the intact dura into the cortex. Neural signals were amplified and processed using an online spike detection and sorting pipeline based on template matching. Templates were defined interactively from initial spike waveforms. Electrode positioning and real-time spike processing were controlled by the AlphaLab system (Alpha Omega Engineering).

## Data analysis

### Neuron selection

Trials were segmented into four epochs during both action observation and execution: approach (from hand release to object contact), grasp (from object contact to movement onset), manipulation (from movement onset to completion of the action), and hold (maintenance of the object at the final position).

We identified neurons that responded during action observation or action execution, following a previously described procedure^16^. For each neuron, responses during the observation and execution phases were evaluated separately. For each of the four conditions, firing rates during the four epochs and the baseline phase (first 500 ms of each trial) were compared using a Friedman test. Tests were performed separately for each condition, and significance was assessed at α = 0.05/4 (Bonferroni-corrected for the four conditions).

A neuron was considered responsive in a given phase (observation or execution) if at least one condition showed a significant effect. Neurons that were responsive in both phases were selected.

### Joint PCA

Our approach is related to demixed PCA but adapted to the block structure of our task, which does not permit a standard ANOVA-style decomposition^10^. Analyses were performed separately for the video-blocked and cue-blocked paradigms. Each paradigm consisted of two blocks (Blocks A and B for the video-blocked paradigm; Blocks C and D for the cue-blocked paradigm), with two conditions per block (see Fig. 2).

For each neuron, spike trains were smoothed using a Gaussian kernel (σ = 50 ms), yielding a continuous firing rate over time. These firing rates were averaged across trials for each condition, resulting in a single time course per neuron and condition. Within each block, firing rates were then soft-normalized by dividing by the maximal firing rate of the neuron in that block plus a constant (max + 5), as in previous work^25^. The constant prevents division by small values and reduces amplification of neurons with low firing rates. For each neuron and time point, the mean firing rate across the two conditions within a block was subtracted, thereby removing activity shared across conditions within that block. This preprocessing ensures that the resulting activity reflects differences between the two conditions that varied across trials within each block (same vs. other in the video-blocked paradigm; lift vs. twist video in the cue-blocked paradigm).

For each paradigm, we denote the two blocks as Block 1 and Block 2 (corresponding to Blocks A and B in the video-blocked paradigm, and Blocks C and D in the cue-blocked paradigm). For a given paradigm and trial phase, the population activity was arranged into data matrices X_1_, X_2_ ∈ ℝ^2T×N^, corresponding to Block 1 and Block 2, respectively. Each matrix was constructed by concatenating the time-resolved activity of all neurons across the two conditions within a block, where T denotes the number of time points per condition and N the number of neurons. For the execution phase, data were aligned to movement onset, and analyses were restricted to the subsequent 1000 ms. Conditions were ordered such that trials corresponding to the same executed action were aligned across blocks.

Given the blocked task structure, the joint PCA was used to identify population components that capture variance shared across blocks and to determine whether this variance was aligned with the executed action or with the task variable that varied across trials in the respective paradigm (rule cue in the video-blocked paradigm; observed action video in the cue-blocked paradigm).

To simplify notation, we describe the procedure for a single trial phase (preparation) in the cue-blocked paradigm. We first consider the principal component ω ∈ ℝ^*N*^ that explains the largest amount of variance in Block 1, defined as

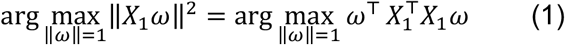

The solution is given by the eigenvector of 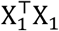 corresponding to the largest eigenvalue. To identify components that maximize variance across both blocks, we extended equation (1) to

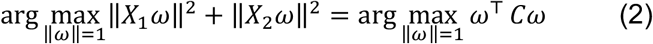

where

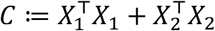

Since *C* is symmetric, its eigendecomposition yields the desired components analogously to the single-block case.

To ensure that components reflect consistent structure across blocks, we additionally required that projections onto the component are similar across blocks. Block 1 contains the conditions

“Lift Video + Lift Action” and “Twist Video + Twist Action”, whereas Block 2 contains “Twist Video + Lift Action” and “Lift Video + Twist Action”. For brevity, we write

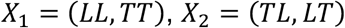

where parentheses denote concatenation of the submatrices. For a given component ω, the projections *X*_1_ω and *X*_2_ω are vectors in ℝ^2T^. To quantify the similarity between projections across blocks, we consider the squared Euclidean distance

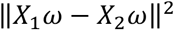

We then expand this expression as

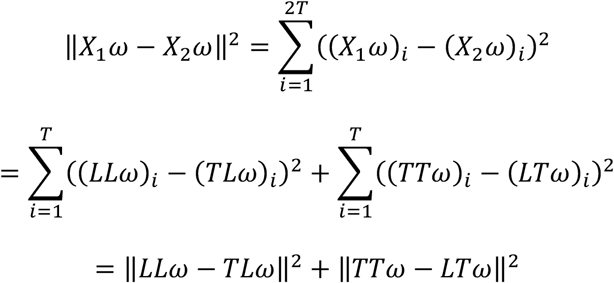

If the distance 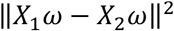 is small, projections corresponding to the same executed action are similar across blocks (LL with TL, TT with LT). Thus, the component ω aligns conditions with identical executed actions, indicating that its variance reflects tuning to the executed action rather than to the observed video.

In contrast, consider the matrix 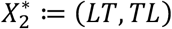, which corresponds to a permutation of the condition order and provides a reference for alignment based on the alternative task variable rather than the executed action. If the distance 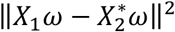 is small, projections align according to the observed action rather than the executed action. Thus, the component ω aligns conditions with identical observed actions, indicating that its variance reflects tuning to the observed video.

Building on equation (2), we derived objective functions for components that maximize variance across both blocks while being selectively tuned to either the executed action or the alternative task variable. The objective for the first action-aligned component is given by

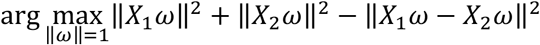

The first video-aligned component is given by

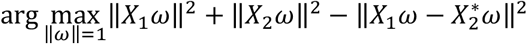

This leads to a unified objective function that does not require prespecifying whether the component is aligned with the executed action or with the alternative task variable. Specifically, components are found by an eigendecomposition that minimizes either of the two partial objective functions above. Writing this as a minimum, we therefore seek the component ω that maximizes

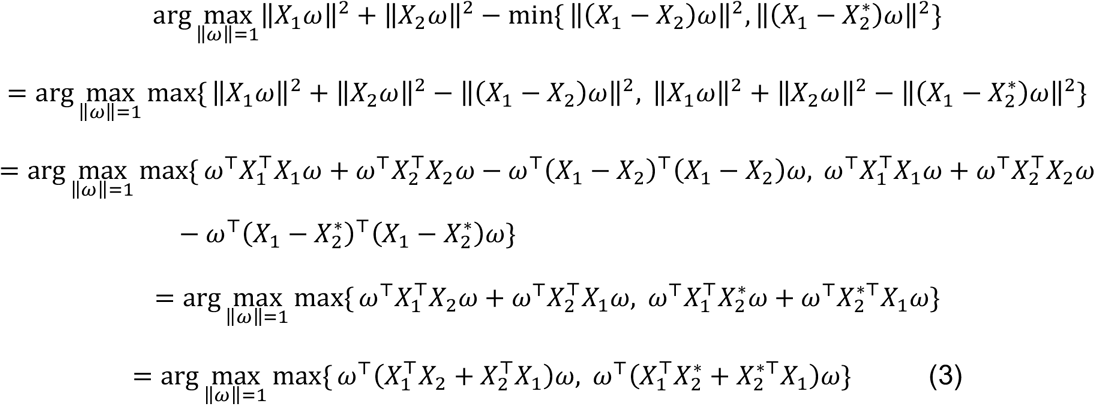

Using 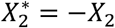, equation (3) simplifies to

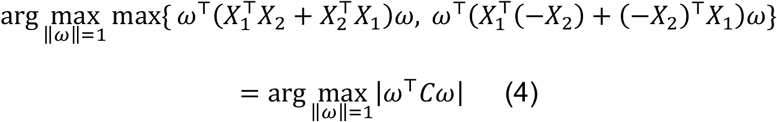

where

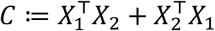

The matrix *C* is symmetric. The eigenvector corresponding to the largest eigenvalue of *C* maximizes the quadratic form ω^T^*C*ω under the constraint ||ω|| = 1, whereas the eigenvector corresponding to the smallest eigenvalue maximizes ω^T^(−*C*)ω. Consequently, the eigenvector associated with the eigenvalue of largest absolute value maximizes equation (4). The sign of the eigenvalue indicates whether the component aligns with the executed action or with the alternative task variable.

The eigenvector corresponding to the eigenvalue with the second-largest absolute value defines the direction that maximizes equation (4) while being orthogonal to the leading eigenvector. Principal components were then obtained by eigendecomposition of *C* and ordering the eigenvectors according to the absolute values of their corresponding eigenvalues. For each component, the sign of the eigenvalue determines whether it is aligned with the executed action or with the alternative task variable.

In practice, we additionally imposed sparsity on the component weights to ensure that the identified components reflected consistent neural contributions across blocks at the single-neuron level. Without this step, neurons active in only one block could still contribute to the joint components, potentially biasing the attribution of variance to different task variables. To mitigate this effect, we penalized non-zero weights for neurons whose activity patterns differed across blocks. After computing each component ω via eigendecomposition as described above, we used ω as the initial value for the following optimization problem

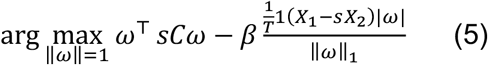

Here, 1 ≔ (1, …,1), and *s* = 1 if the component is aligned with the executed action and *s* = −1 otherwise. Thus, *s* flips the sign of *C* and *X*_2_, allowing both alignment conditions to be expressed within a single objective.

Compared to equation (4), the variance-maximizing term remains unchanged. We additionally introduced an *L*_1_-penalty on ω, such that dimensions in which *X*_1_ and *X*_2_ differed more strongly are penalized more heavily, resulting in a weight vector that relies on dimensions shared by *X*_1_ and *X*_2_.

Although the resulting objective is not convex, initializing the optimization at the analytical solution of the non-sparsified problem consistently yielded stable components. For each component, we selected the smallest value of β such that the regularization term remained below the average across neurons during the baseline period. This procedure reduced contributions from neurons with inconsistent activity across blocks, except for differences attributable to noise.

### Generalized linear model analysis

To assess whether single-neuron responses were modulated by the rule and the required self-action beyond the observed action, we fitted generalized linear models (GLMs) separately for each neuron and for each trial phase (cue and observation) in the video-blocked paradigm. Neuronal responses were quantified as spike counts within the respective analysis window. Spike counts were modeled using a Poisson distribution.

In the video-blocked paradigm, the observed action (video) was constant within each block, whereas the rule (same vs. other-action) varied across trials. This design allowed us to examine how neuronal responses to the same observed action depended on the rule and on the required self-action. For each neuron, we fitted models in which spike counts were expressed as a function of the rule, the required self-action (lift vs. twist), and their interaction. The required self-action was determined by the combination of observed action and rule. To determine whether neuronal responses were better explained by the observed action or by the rule and the required self-action, we compared a model based solely on the observed action with a model based on the rule, the required self-action, and their interaction. Model comparison was performed using a likelihood-ratio test. Neurons for which the model including rule, required self-action, and their interaction provided a significantly better fit were classified as exhibiting modulation beyond the observed action and included in subsequent analyses.

To characterize the nature of task-related modulation, we examined the regression coefficients associated with self-action (*β*_*A*_) and the interaction between observed action and self-action (*β*_*AV*_). To facilitate interpretation of the regression coefficients, observed action was coded as a binary variable and required self-action as a binary variable with opposite signs for the two actions, such that specific relationships between the coefficients correspond to distinct tuning profiles. Intuitively, *β*_*A*_ captures how strongly a neuron differentiates between the two self-actions, while *β*_*AV*_ reflects whether this difference depends on the observed action. If *β*_*AV*_ ≈ 0, the effect of self-action is the same for both observed actions, indicating tuning to the required self-action independent of the observed video. If *β*_*AV*_ ≈ −*β*_*A*_, the self-action effect is present for one observed action but absent for the other, indicating selectivity for a specific combination of observed action and required self-action. Finally, if *β*_*AV*_ ≈ −2*β*_*A*_, the self-action effect reverses depending on the observed action, such that neuronal responses distinguish between same and other conditions rather than between lift and twist. This pattern corresponds to tuning to the rule, which determines how the observed action is mapped onto the required self-action. To assign neurons to these categories, we computed the distance of the observed coefficient pair (*β*_*A*_, *β*_*AV*_) to these three idealized cases and classified each neuron according to the closest match.

